# Hopeful monsters: Unintended sequencing of famously malformed mite mitochondrial tRNAs reveals widespread expression and processing of sense-antisense pairs

**DOI:** 10.1101/2020.09.08.286963

**Authors:** Jessica M. Warren, Daniel B. Sloan

## Abstract

Although tRNA structure is one of the most conserved and recognizable shapes in molecular biology, aberrant tRNAs are frequently found in the mitochondrial genomes of metazoans. The structure of several mitochondrial tRNAs is so degenerate that doubts have been raised about their expression and function. Mites from the arachnid superorder Acariformes are predicted to have some of the shortest mitochondrial tRNAs, with apparent base mismatches in acceptor stems and a complete loss of cloverleaf-life shape. While performing mitochondrial isolations and recently developed RNA-seq methods to capture mature, CCA-tailed tRNAs in plant tissue, we inadvertently sequenced the mitochondrial tRNAs from a common plant pest, the acariform mite *Tetranychus urticae,* to a high enough coverage to detect all previously annotated *T. urticae* tRNA regions. The results not only confirm expression, CCA-tailing and post-transcriptional base modification of these highly divergent tRNAs, but also revealed widespread sense and antisense expression of tRNA genes in the *T. urticae* mitochondrial genome. Mirrored expression of mitochondrial tRNA genes has been previously hypothesized but not demonstrated to be extensive in any system. We discuss the functional roles that these divergent tRNAs could have as both decoding molecules in translation and processing signals in transcript maturation pathways for other genes, as well as how the surprising finding of sense-antisense tRNA pairs adds another dimension to the bizarre tRNA biology of mitochondrial genomes.

## Introduction

As the adapter molecule between mRNA codons and amino acids, transfer RNAs (tRNAs) are a conserved feature of life. The vast majority of known tRNAs have a uniform structure comprised of acceptor and anticodon stem, D- and T-arms, a variable region, and accompanying loops to form the easily recognizable clover-leaf secondary structure. An increasing number of exceptions to this structural uniformity has come from the expanding field of mitochondrial genetics. Most animal mitochondrial genomes encode 22 tRNA genes considered sufficient to decode all codons [1, 2] (but see [3, 4]). However, some mitochondrial tRNAs (mt-tRNAs) can deviate significantly from the canonical tRNA structure. First identified in mammalian mitochondria in the late 1970s, a tRNA-Ser lacking the D-arm has now been found to be very common in metazoan mitochondria [5–8]. Following the mt-tRNA-Ser discovery, even more extensive cases of tRNA truncation were reported in nematode mitochondria where all of the mt-tRNAs lacked either the D- or the T-arm, which were lost in favor of short replacement loops consisting of only five to eight nucleotides [9, 10]. The most extreme examples of tRNA truncation have come from a class of nematodes (Enoplea) and a superorder of arachnids (the acariform mites)[11–13]. Aside from the mammalian mitochondrial tRNA-Ser lacking the D-arm, there is very limited experimental evidence for expression or aminoacylation activity in translation of these aberrant tRNA structures [2, 4]. The most notable support comes from the sequencing of several mt-tRNAs from the nematode *Romanomermis culicivorax,* including an armless tRNA that was shown to be post-transcriptionally modified with a CCA-tail [11]. In the Acariformes, mt-tRNA expression data are entirely lacking, and unlike in Enoplea, mt-tRNAs in Acariformes are not well conserved, with some tRNA genes differing even at an intraspecific level [13]. The highly divergent and aberrant structure of mt-tRNAs in acariform mites poses questions about the limits of what constitutes a minimal functional tRNA [12, 14, 15].

Mites are among the oldest and most diverse groups of terrestrial animals [16]. As one of the two superorders of mites, the Acariformes include well-known representatives such as dust mites, scabies, chiggers, and spider mites. Partially due to both agricultural and medical concerns, there has been increasing interest in the mitochondrial genomics of acariform mites for phylogenetics as well as development of pest control agents and treatments [15, 17]. One of the most notable features discovered from the sequencing of Acariformes mitochondrial genomes has been the widespread rearrangement and extreme truncation of mt-tRNAs [13, 14, 18–20]. Going beyond the frequently described loss of a D- or T-arm, mt-tRNAs from multiple taxa in Acariformes have been predicted to completely lack both arms and produce just a simple stem-loop structure [13]. In other species, no structure identifiable as a tRNA could be found for some anticodons, and they were reported as functionally lost [12, 18, 21, 22]. In addition to their extremely short lengths, some predicted acariform mt-tRNAs have mismatches in acceptor stems [12, 13, 20], which appear to be widespread in diverse arachnid lineages and often occur at the interface of overlapping tRNA genes [23, 24]. It is possible that these mismatches are edited and corrected post-transcriptionally as observed in some other eukaryotes [25–27]. Regardless of whether they are edited, these mismatches create additional difficulties when trying to annotate tRNA genes and predict structure and function.

Programs like tRNAscan-SE [28] and ARWEN [29] that are used to infer tRNA gene presence from genomic data frequently fail to detect the highly degenerate mt-tRNAs in Acariformes taxa. The true loss of a mt-tRNA gene would likely require that a nuclear-encoded tRNA be imported into the mitochondrial matrix to as a functional replacement [30] – a phenomenon that is ubiquitous in some eukaryotic lineages (e.g. plants [4, 31]) but thought to be rare in metazoans [2]. Doubts about the extensive import of tRNAs into animal mitochondria have led to the revisiting and manual inspection of noncoding space in multiple Acariformes mitochondrial genomes, resulting in predictions for additional, but sometimes not all, mt-tRNA genes [12, 13, 15, 22].

An additional complication of proposed mt-tRNA losses relates to the multi-functionality of tRNAs in mitochondrial gene expression. Mt-tRNAs have long been thought to be processing signals for mRNA and rRNA transcript maturation. Under what is known as the tRNA punctuation model, protein-coding and rRNA genes are often flanked by one or more tRNA genes in mitochondrial genomes, effectively co-opting the recognition and cleavage by the tRNA-interacting enzymes RNase P and RNAse Z to produce excised mRNA or rRNAs from polycistronic mitochondrial transcripts [32, 33]. Mt-tRNAs have also been implicated as origins of mitochondrial DNA replication, where hybridization between the tRNA and the complementary DNA gene sequence initiates replication [34, 35]. Thus, the complete loss of some mt-tRNA genes may necessitate additional evolutionary mechanisms to maintain functional mitochondrial gene expression.

Extensive work had been done to predict tRNA candidates as well as assign functionality solely based on genomic data in Acariformes mites [12], but there remains a lack of experimental data for their expression and processing. This scarcity of expression data may be partly attributed to longstanding technical difficulties in sequencing tRNAs (tRNA-seq), due to tRNAs being incredibly recalcitrant to reverse transcription [36], a step necessary in the vast majority of high-throughput RNA-seq methods. tRNAs are the most extensively modified RNAs known [37], and chemical modifications at the Watson-Crick face of tRNA bases can interfere with reverse transcriptase activity. The stalling, skipping, or disassociation of a reverse transcriptase can result in termination of cDNA synthesis or the misincorporation of incorrect nucleotides in the cDNA at the corresponding modified base positions [38–40]. Additionally, the tightly base-paired 3′- and 5′-termini of tRNAs can inhibit adapter ligation, which is necessary for the priming of reverse transcription in RNA-seq library preparation [41]. A major breakthrough in tRNA-seq came from the application of the demethylating enzyme AlkB to remove certain reverse transcription-inhibiting modifications prior to tRNA-seq library construction [42, 43]. Additional improvements in the sequencing of full-length tRNA molecules included the development of adapters that take advantage of the unique structure of mature tRNAs [44, 45]. All mature tRNAs have protruding, unpaired bases at the 3′-termini, composed of the discriminator base and the trinucleotide CCA [46]. This CCA “tail” is post-transcriptionally added by tRNA nucleotidyltransferases and has been considered a hallmark of tRNA functionality as the site of aminoacylation [47]. YAMAT-seq is a recently developed technique based on adapters that are complementary to these protruding nucleotides, aiding adapter ligation and resulting in the capture and sequencing of full-length, mature tRNAs [45].

Here, we report the sequencing of the mt-tRNA complement from the acariform spider mite and common plant pest *Tetranychus urticae*, which was the inadvertent outcome of applying mitochondrial enrichment techniques and targeted tRNA-seq methods to infested plant tissue. This fortuitus capture of the famously divergent mt-tRNAs from *T. urticae* provides unprecedented insight into the expression and post-transcriptional modifications of some of the most aberrant tRNAs ever found.

## Results

### The highly degenerate mt-tRNA complement of *T. urticae* is expressed and post-transcriptionally processed

The application of a gradient-based mitochondrial isolation method on leaf tissue from the angiosperm *Silene vulgaris* resulted in the co-enrichment of the mitochondria from common plant pests inhabiting the tissue alongside the intended plant mitochondria. When these mitochondrial-enriched fractions were subjected to targeted tRNA-seq methods that preferentially capture mature tRNAs (three biological replicates processed both with and without AlkB enzymatic treatment), they produced a substantial number of reads mapping to the mt-tRNA genes of the spider mite *T. urticae*. Reads mapping to all previously annotated *T. urticae* mitochondrial tRNA gene regions were sequenced in all three AlkB-treated libraries and in two of three untreated libraries (only tRNA-Gln, tRNA-Leu2, and tRNA-Trp were not detected in the other untreated library; supp. Table 1). Similar to other tRNA expression profiles described utilizing YAMAT-seq [45, 48], there was a high degree of variability in coverage, with the abundance of the most and least frequently sequenced tRNAs spanning more than three orders of magnitude (Fig. 1, supp. Table 1). Approximately 97% of reads mapping to a tRNA reference contained a 3′ CCA tail, which is typical of the mature, processed tRNAs that are targeted by YAMAT-seq (supp. Table 2).

**Fig. 1.**
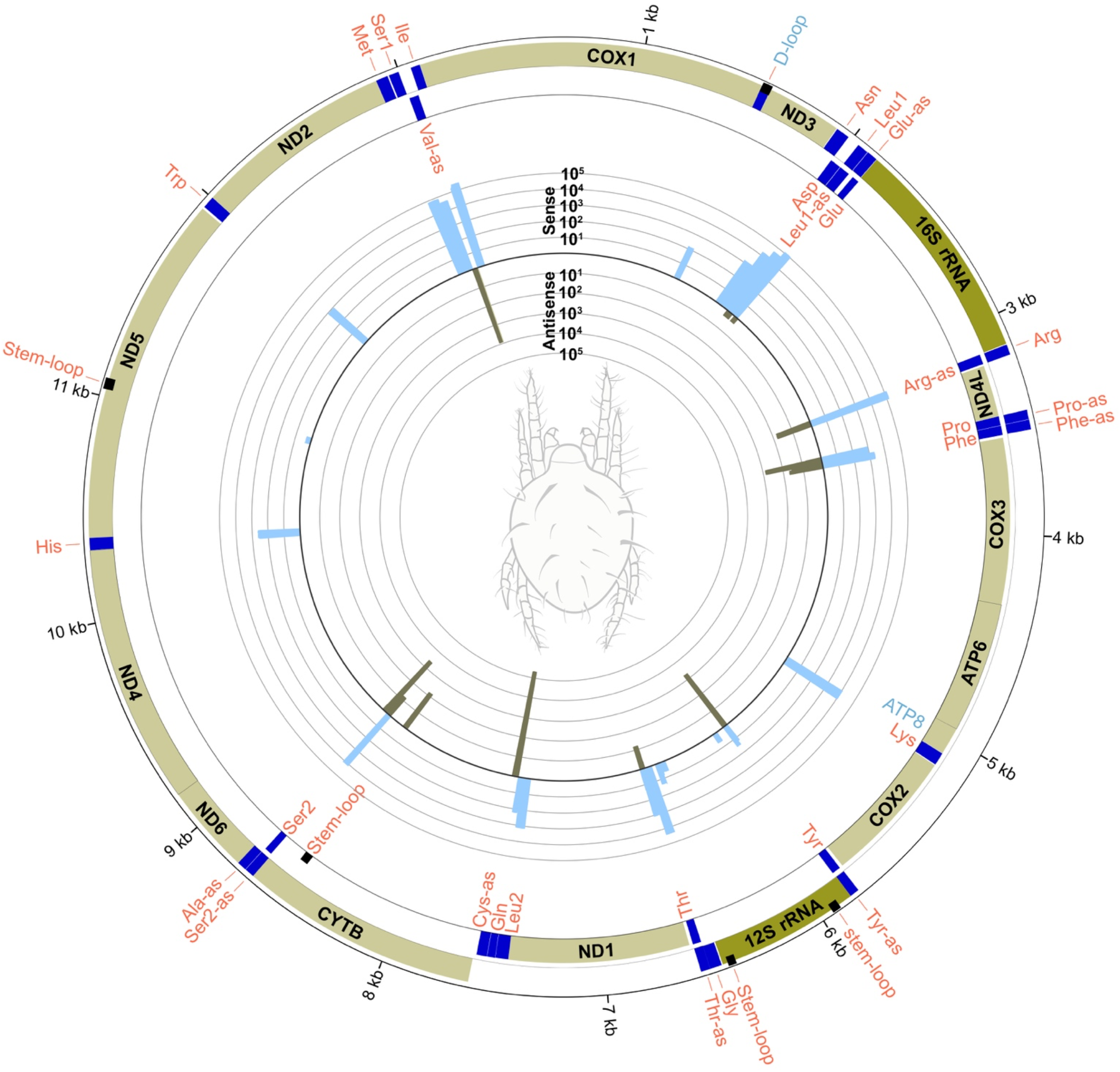
Mitochondrial genome map of *T. urticae* with tRNA and stem-loop expression. Outer circle shows the gene organization including expressed and CCA-tailed sequences determined by this study. Light green blocks are protein coding genes, dark green are ribosomal RNAs, black are stem-loops, and dark blue are mt-tRNAs. The two gene tracks represent gene transcribed on the majority strand (outer track) and minority strand (inner track). The histograms show the sense (outward-facing, light blue bars) and antisense (inward-facing, dark gray bars) detection of reads mapping to each tRNA or stem-loop on a log_10_ scale. Sense and antisense expression are defined relative to prior *T. urticae* mitochondrial annotations [13]. Diagram was created using Circos (ver. 0.69-9)[74].

The majority of transcripts corresponded to the length and start/stop coordinates predicted from the prior annotation of the sequenced *T. urticae* mitochondrial genome [13] with some minor truncations compared to the genomic annotations; however, a few tRNAs appear to deviate considerably from the predicted models. tRNA-Phe and tRNA-Thr were previously annotated as having a D-arm, but all transcripts mapping to these genes were considerably shorter, completely lacking both arms (Fig. 2, supp. Table 2).

**Fig. 2.**
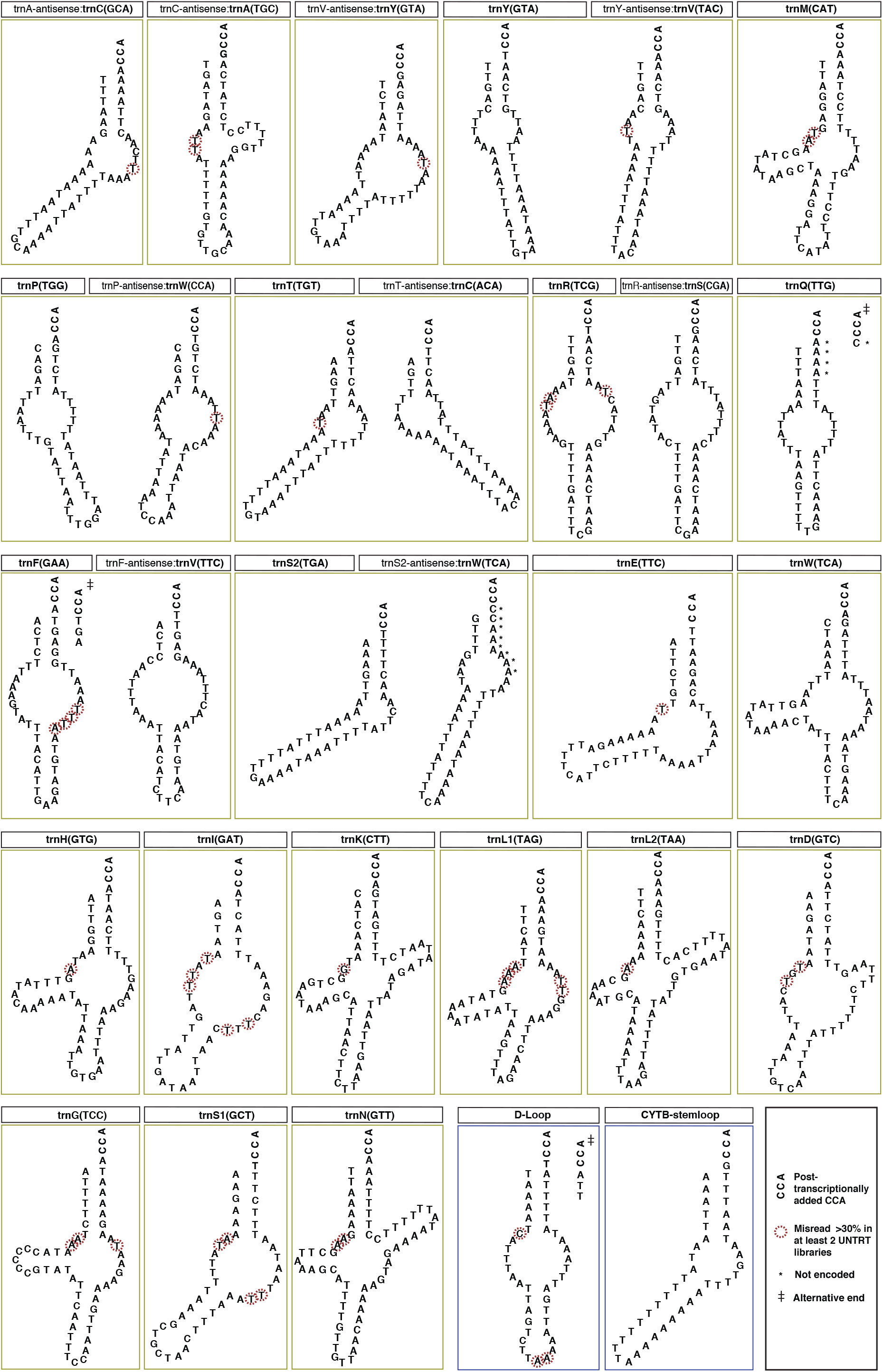
Predicted secondary structures of *T. urticae* mt-tRNAs and stem-loops. Most structures presented represent the maximum free energy folding model using the program the program RNAfold [73] (see methods for modification of folding predictions for some tRNAs). The primary reference sequence is based on the most highly detected YAMAT read mapped to the reference that had 100% base identify to the genomic sequence. Bases were inferred to be modified if >30% of the reads at a position differed from the reference sequence in untreated libraries and are highlighted with a dashed red circle. Bases detected in sequenced reads that were not present in the genomic reference are indicated with an asterisk. Some tRNAs had frequently occurring alternative 3′- and 5′-ends which are presented next to the structure and indicated with a ‡. Folded structure diagrams were created using the program VARNA (ver. 3.9, [75]).

More surprisingly, the template strand of multiple tRNAs was found to be opposite to what has been previously annotated. Many RNA-seq methods do not retain the strand information of the original RNA template due to the synthesis and adapter ligation of randomly fragmented and primed double-stranded cDNA molecules [49]. However, annealing of Y-shaped YAMAT-seq adapters directly to tRNA molecules in a defined orientation (as well as the presence of the CCA tail) provides 5′- and 3′-prime information, allowing for the identification of the transcribed strand [45]. Strand information is particularly useful in tRNA-seq because the structure of tRNAs as well as the complementary nature of nucleic acid base-pairing means that the reverse complement of a tRNA gene can often produce a similar, “mirrored” folding structure. Therefore, both strands could be predicted to produce a functional tRNA, albeit with different anticodons. Without experimental evidence and conservation of tRNA arrangement in a mitochondrial genome, inferring a tRNA’s identity and sense-orientation from genomic data alone can require a process of elimination searching for anticodons, and in cases of “reciprocal pairs” of anticodons that are complementary to each other, some guesswork. For example, in the dust mite *Dermatophagoides pteronyssinus*, the same region of the mitochondrial genome has been annotated to encode either tRNA-Val or its antisense complement tRNA-Tyr depending on the study [12, 19], demonstrating the challenge in assigning tRNA identities to genomic sequences in which complementary expression can also form tRNA-like structures. In the case of *T. urticae*, no sense expression of tRNA-Cys, tRNA-Val, or tRNA-Ala was detected (Fig. 1). Note that tRNA-Cys and tRNA-Ala have complementary anticodons (GCA and TGC, respectively). Therefore, the finding of exclusive antisense expression for both of these two genes effectively “switches” the anticodon/tRNA identity of the genes relative to their annotations in the genome. Similarly, the antisense-only expression of the tRNA-Val could be functionally compensated by the predominantly antisense expression of tRNA-Tyr (Fig. 2), as these tRNAs also have complementary anticodons (GTA/TAC).

### Expression and processing of non-tRNA stem-loops

Although the vast majority of transcripts mapped to a predicted tRNA gene, there was a detectable level of reads originating from specific positions within 12S rRNA, the putative control region (D-loop), a region internal to *ND5*, and an antisense region internal to *CYTB*, all with a post-transcriptionally-added CCA-tail (supp. Table 2 and Fig.1). Post-transcriptional addition of CCA to non-tRNAs has now been described in the mitochondria of numerous other species, including rat [50], human [51], and plants [52]. All of the apparent non-tRNAs that we detected in *T. urticae* have a predicted stem-loop structure (supp. Table 3), which may make them effective substrates for tRNA-processing machinery.

### The novel detection of sense-antisense expression of tRNAs

In addition to cases where only antisense reads were found (see above), mirrored expression of both sense and antisense transcripts was also detected for a number of *T. urticae* mt-tRNA genes. Specifically, tRNA-Arg, tRNA-Glu, tRNA-Leu1, tRNA-Phe, tRNA-Pro, tRNA-Ser2, and tRNA-Tyr all had both sense and antisense reads mapped but with variable percentages (Fig. 1, supp. Table 2). Very little sense expression of tRNA-Tyr was detected, with over 98% of all the reads mapped to the reference being in the antisense direction with an anticodon of TAC (Val). Conversely, tRNA-Glu and tRNA-Leu1 had only a single antisense read sequenced and thus lack evidence for a meaningful level of antisense expression. The remaining tRNAs with both sense and antisense expression all had at least seven (and up to 348) antisense reads, representing anywhere from <1% to 43% of the mapped sequences the respective reference gene.

To assess whether the mirrored sense-antisense expression observed for *T. urticae* mt-tRNAs was a general phenomenon or perhaps an artefact of the tRNA-seq method, we examined the abundant mt-tRNA reads generated from the intended host plant species, *S. vulgaris*. We found that a total of five (imperfectly matched) reads were generated from a single plant mitochondrial gene (mt-tRNA-Trp) out of 105,867 reads that were mapped to that tRNA (representing 0.00005%). There were no antisense reads and 1,215,689 sense reads for the remaining four *S. vulgaris* mt-tRNA reference genes. We also analyzed reads derived from *S. vulgaris* plastids and found that there was a total of just 39 reads (or 0.000014% out of 2,532,488 total mapped for the 30 plastid tRNA genes) that could have an antisense orientation, and many of these reads failed a hit coverage threshold in mapping (i.e. fragments). Thus, the abundant sense-antisense expression appears to be a distinct feature of *T. urticae* mt-tRNAs and not a universal outcome of tRNA-seq in these heterogeneous samples. However, caution should be applied when interpreting the relative expression between sense and antisense transcripts from the same gene, as substantial reverse-transcription biases still exist in tRNA-seq datasets [45, 48]. Similarly, the lack of detection of a tRNA transcripts, in either orientation, should not be taken as definitive evidence of lack of expression.

### Evidence for base modifications affecting reverse transcription at positions 8 and 9 and 3′-end processing of overlapping tRNAs

Misincorporation of incorrect nucleotides during the reversion transcription of tRNAs has been used to predict the modification state of tRNA bases [39, 40]. We considered a base confidently modified if it was misread in >30% of the reads from untreated libraries (Fig. 2). No position had a 100% misincorporation rate, but position percentages ranged up to 99% misread in all libraries (position 9T in antisense tRNA-Cys; supp. Table 4). Thymines at positions 8 and 9 and adenines at position 9 in *T. urticae* mt-tRNAs were the most likely to exhibit differences relative to the reference sequence (Fig. 2, supp. Table 4, supp. Fig. 1). The percentage of reads with the correct base at position 9A greatly increased in at least two of the three AlkB-treated libraries for tRNA-Asn, tRNA-His, tRNA-Leu2, and tRNA-Met (supp. Table 4, supp. Fig. 1). AlkB treatment did not appear to be as effective in one of the replicates (replicate 2). An apparent error in quantifying RNA concentration during AlkB treatment of replicate 2 resulted in a higher ratio of RNA to AlkB and may account for some replicate behavior. The misincorporation signal was often almost eliminated (i.e. the correct base was read) after AlkB treatment in replicates 1 and 3 for position 9A in tRNA-Asn, tRNA-His, tRNA-Leu2, and tRNA-Met, whereas it was only reduced (but not eliminated) in replicate 2 (supp. Table 4, supp. Fig. 1). The most frequently misread base was a 9T, and the number of reads with the correct base did not increase after AlkB treatment (i.e. an apparently AlkB-insensitive modification) (supp. Fig. 1, supp. Table 4).

Some transcripts also contained nucleotides (adenosines and cytosines in addition to the 3′ CCA tail) not encoded in the mitochondrial genome, including sense reads from tRNA-Gln and antisense reads from tRNA-Ser2 (Fig. 2), which suggests that these nucleotides may be post-transcriptionally added. Prior tRNA gene annotations for tRNA-Gln had predicted a mispaired 4A:43C and overlap with tRNA-Leu2 [13]. However, sequenced reads suggest that transcription of genomically encoded sequence ends at position 38 (supp. Table 2). Similarly, the antisense expression of tRNA-Ser2 appears to end at position 36 and is finished with the post-transcriptional addition of nucleotides, thereby preventing overlap with the antisense expression of tRNA-Ala (Fig. 2, supp. Table 2).

## Discussion

The predicted sequence and structure of aberrant mt-tRNAs in multiple metazoan systems has raised questions about their expression, maturation, and functionality because of short sequence length, base mispairing in stems, and a lack of canonical tRNA structure [5, 9, 53–55]. These extreme deviations from canonical tRNA structure have made identifying mt-tRNA with genomic information alone difficult, leading to some studies inferring their outright loss [14, 18, 21, 22]. Our accidental sequencing of *T. urticae* mt-tRNAs, an arachnid from a superorder (Acariformes) well known for degenerate mt-tRNAs, has confirmed that previously predicted tRNA genes, some as short as 44 nt at maturity, are transcribed and modified with the post-transcriptional addition of the CCA-tail (supp. Table 2). There is also evidence from sequencing misincorporations at specific sites (especially positions 8 and 9) that these tRNAs have post-transcriptionally modified bases. One position, 9A, was frequently misread in T-armless mt-tRNAs and appeared to be removed by AlkB treatment (supp. Fig 1, supp. Table 4). One possibility is that this site carries a 1-methyladenosine (m^1^A_9_) modification, which is known to be removed by AlkB [39, 42, 43] and has been found to be necessary for efficient aminoacylation and EF-Tu-binding of T-armless mt-tRNAs in nematodes [56]. Misincorporations at positions 8T and 9T were most common, and we hypothesize that these positions are modified to prevent base pairing with A/T rich replacement loops, thereby maintaining the L or boomerang-like structure necessary for ribosome interaction [11, 57]. We advise that any specific sites of interest should be investigated for possible contributions of sequence variation at the genomic level to apparent misincorporation patterns. Overall, the expression, end-processing, and base modification of *T. urticae* mt-tRNAs suggest that the mt-tRNA gene complement is functional and sufficient to decode the mitochondrial genome despite extremely degenerate structures.

However, caution is warranted when inferring a role in translation because stem-loop structures (e.g. in the D-loop and certain non-tRNA genes) were also expressed and processed, and it is unlikely that these function in translation as tRNAs. Although the post-transcriptional modification of tRNAs and especially the addition of the CCA-tail have been frequently used as evidence for functionality of tRNAs [10, 11], the detected processing of apparent non-tRNA stem-loops in *T. urticae* and other systems [50–52] may mean that the addition of a CCA-tail alone is insufficient to assign function as a tRNA in translation. Aminoacylation and EF-Tu binding assays will be necessary to determine whether tRNAs and stem-loop structures are functional in translation or have other functions in the organelle.

Mt-tRNAs have long been hypothesized to have additional functional roles in both vertebrate and invertebrate mitochondrial expression based on co-opting tRNA-interacting enzymes (RNase P and Z) for mRNA and rRNA transcript maturation through the endonucleolytic cleavage of polycistronic RNAs, releasing transcripts immediately before and after a tRNA sequence [32, 58]. The sense and antisense expression of mitochondrial tRNA-Tyr and tRNA-Arg in *T. urticae* occur at protein-coding boundaries on both strands, raising the possibility that tRNA genes can act in a novel form of dual-strand punctuation by being recognized by enzymes in both orientations. As such, the conservation and expression of tRNAs or tRNA-like structures in acariform mitochondria could be for the maintenance of a structure generally recognizable by enzymatic partners in transcript maturation pathways for other genes [19], as well as possible replication initiation sites [34, 35]. The loss of translation-related function of a mt-tRNA but the conservation of some tRNA structural features (i.e. the acceptor stem) for transcript maturation has been suggested for marsupial mt-tRNA-Lys, where a nonfunctional tRNA-Lys is retained in the mitochondrial genome as a processing signal despite its functional replacement by an imported nuclear-encoded tRNA-Lys [59]. However, the antisense expression and modification of multiple mt-tRNAs in *T. urticae* not immediately up or downstream of any previously annotated genes suggests that expression may be maintained for functions other than as a processing signal. Alternatively, some processed sequences may simply be a byproduct of being in-between two sense expressed tRNAs. In addition, there appears to be extensive modification of some tRNAs, including potential post-transcriptional 3′ terminus “finishing” through the addition of nucleotides other than the CCA-tail (Fig. 2). In the case of tRNA-Gln, these nucleotide additions appear to restore a correctly base paired acceptor stem when a base mispairing exists in the genomic sequence. Taken together, the extensive 3′-terminus modification and possibly stabilizing chemical base modifications (i.e. positions 8 and 9) do suggest the conservation of pathways to maintain structure for enzymatic and ribosome interaction.

If all or most of the *T. urticae* mt-tRNAs are functional in translation, the structural variability of the mt-tRNAs pool raises additional questions about the enzymatic partners necessary for mt-tRNA function. Enzymes and complexes that interact with tRNAs (e.g. EF-Tu and the ribosome) often use conserved structural features such as the T-arms for substrate recognition and interaction [60, 61]. The coexistence of tRNAs lacking either or both arms in addition to tRNAs with conventional shape (i.e. both arms present) results in *T*. *urticae* having a mt-tRNA pool with extreme structural variability. A similar situation in nematodes has arisen where mt-tRNAs either lack a D- or T-arm such that tRNA interactions require two distinct EF-Tu enzymes, one binding only to T-armless tRNAs and the other binding to D-armless Ser-tRNAs [62, 63]. What coevolutionary features of nuclear-encoded enzymes facilitate both degenerate as well as variable mt-tRNA structure is Acariformes is yet to be investigated. Given that highly degenerate mt-tRNAs have arisen independently in multiple lineages, the evolution of the tRNA-interacting enzymes may offer clues into the propensity for the evolution divergent mt-tRNAs [53, 61].

The surprising detection of sense-antisense expression of multiple tRNAs theoretically expands the function of *T. urticae* mt-tRNAs as decoding molecules in mitochondrial protein synthesis. The possibility of a single mt-tRNA gene serving as two different isoacceptors has been suggested for the nematode *Romanomermis iyengari* where tRNA-Ala and tRNA-Cys are predicted as superimposed reverse complements at the same genomic location [51], and other reports have found some limited evidence of antisense mt-tRNA expression [64, 65]. However, dual sense-antisense expression of mt-tRNA genes has remained a largely undetected phenomenon despite the obvious potential for sense and antisense transcripts to produce mirrored secondary structures [66]. Interestingly, sense-antisense expression of mt-tRNAs has been proposed to account for the highly pathogenic nature of mutations in mt-tRNA genes [66]. Disease causing mutations are 6.5 times more frequent in mt-tRNAs than in other mitochondrial genes and this discrepancy in pathology may arise from tRNAs having multiple functions, including dual roles in translation as two different tRNAs. The complementary expression of tRNAs has even been suggested to represent the ancestral decoding system [67]. The transcription of two different tRNAs from a single tRNA gene could also facilitate mitochondrial genome reductions through the tolerance of mt-tRNA loss. As such, our findings in support of the long-hypothesized potential for mirrored sense-antisense expression of tRNAs has broad implications for the function and evolution of mitochondrial genomes.

## Materials and methods

### Mitochondrial isolation, RNA extraction, AlkB treatment and YAMAT-seq

Mitochondrial isolations were performed on *T. urticae* leaf tissue from the angiosperm *S. vulgaris*. Plants were grown in a greenhouse with supplemental lighting (16-hr/8-hr light/dark cycle) in the Colorado State University greenhouse, Fort Collins, CO. All mitochondrial isolations were done in triplicate. Each replicate used 75 g of leaf tissue from a dozen 8-month old plants. Leaf tissue was disrupted in a Nutri Ninja Blender for 2 x 2-sec short bursts, and 1 x 4-sec blending in 350 mL of a homogenization buffer containing: 0.3 M sucrose, 5 mM tetrasodium pyrophosphate, 2 mM EDTA, 10 mM KH_2_PO_4_, 1% PVP-40, 1% BSA, 20 mM ascorbic acid, and 5 mM cysteine, pH 7.5-KOH.

Differential centrifugation was performed to remove nuclei, plastids, and cellular debris in a Beckman Avanti JXN-30 centrifuge with a JS-24.38 swinging bucket rotor with the following centrifugation steps: 1) 10 min at 500 g with max brake, 2) 10 min at 1500 g with max brake, 3) 10 min at 3000 g with max brake. After each centrifugation step, the supernatant was transferred into a clean centrifuge tube. Mitochondria were then pelleted by centrifugation for 10 min at 20,000 g with the brake off. Supernatant was discarded and the pellet was resuspended using a goat-hair paintbrush and 2 mL wash buffer containing 0.3 M sucrose, 10 mM MOPS, 1 mM EGTA, pH 7.2-KOH. 30 mL wash buffer was then added to the resuspended pellet and the homogenate was centrifuged for 5 min at 3000 g. Mitochondria were once again pelleted by centrifugation for 10 min at 20,000 g. The pellet was resuspended with 500 *μ*L wash buffer and paint brush. The mitochondrial supernatant was then added to a glass Dounce homogenizer and homogenized with three strokes.

Homogenized mitochondria were then suspended on top of a Percoll gradient with the following Percoll density layers, 18%, 25%, 50%. The gradient was then centrifuged at 40,000 g for 45 min with the brake off. The mitochondrial band at the 25%:50% interface was then aspirated off of the gradient and diluted with 30 mL of wash buffer. The diluted mitochondria were then centrifuged at 20,000 g for 10 min. The supernatant was vacuum aspirated, and the mitochondrial pellet was resuspended in a fresh 30 mL of wash buffer and centrifuged at 10,000 g for 10 min. The supernatant was vacuum aspirated, and the mitochondrial pellet was resuspended in a fresh 1000 *μ*L wash buffer. Resuspended mitochondria were centrifuged at 10,000 g for 10 min. Supernatant was removed with a pipette and the mitochondrial pellet immediately went into RNA extraction procedures. RNA isolations, AlkB treatment, YAMAT-seq library construction, and sequencing were performed using previously published methods [48]. Sequencing reads are available via the NCBI Sequence Read Archive under BioProject PRJNA662108.

### YAMAT-seq read processing and mapping

Reads were trimmed with Cutadapt (ver.1.16 [68]) using the following options: -q 10 --discard-untrimmed --nextseq-trim=20. Forward and reverse trimmed reads were merged using BBMerge [69] (BBTool software package), with a minimum overlap of 20 bp and 0 mismatches. Identical reads were summed and collapsed into read families using the FASTQ/A Collapser tool from the FASTX-Toolkit version 0.0.13 (http://hannonlab.cshl.edu/fastx_toolkit/index.html). The mapping of reads to reference tRNA gene set (described below) was done with previously published pipeline and Perl scripts ([48], https://github.com/warrenjessica/YAMAT-scripts). The number of reads with a CCA tail was also calculated with a custom Perl script. Modified sites that differed from the reference were determined with a pre-existing pipeline [48], except MUSCLE (ver. 3.8.31, [70]) was used as the alignment tool. Pipeline scripts can be found at https://github.com/warrenjessica/YAMAT-scripts.

### Extraction and taxonomic identification of *Tetranychus* reads

While applying a pipeline to remove commonly occurring environmental contaminants (e.g. soil bacteria) it became apparent that a considerable number of reads mapped to *Tetranychus* mitochondrial genome accessions deposited on GenBank. Preliminary analysis found that *T. urticae* (Genbank accession KJ729022) was the most common species. In order to extract all possible reads originating from the *T. urticae,* reads were BLASTed (blastn, e-value of < 1e-3, low complexity regions not filtered [-dust no]) against the *T. urticae* mitochondrial genome and only those that produced a hit were retained for further filtering. To remove any tRNAs originating from other organisms, reads that hit to the *T. urticae* mitochondrial genome were then BLASTed against the GenBank nt nucleotide database (downloaded 1/28/2020), and the taxonomic hit information for the top hit for each read was extracted. Only reads in which the top hit had a taxonomic assignment containing the genus *Tetranychus* were retained. The entire mitochondrial genome of *T. urticae* has been included as a contaminating contig in the whole genome assembly of the flowering plant *Dioscorea rotundata* (Genbank accession LC219377). Therefore, reads with a top hit to this *D. rotundata* accession were also retained.

To determine the *Teranychus* species and strain that was sequenced, the filtered *Tetranychus* reads were mapped to the eight NCBI RefSeq *Tetranychus* reference mitochondrial genomes (HM753535, KJ729017, KJ729018, KJ729019, KJ729020, KJ729021, KM111296 and NC_010526) as well the green strain of *T. urticae* (KJ729022) and the red strain of *T. urticae* (KJ729023) published by Chen et al., 2014 [13] Of these reads, the overwhelming majority (94%) showed a best hit (or a tied best it) to *T. urticae* strain green (KJ729022). The small minority of reads that had a better match to another *Tetranychus* species or strain of *T. urticae* (i.e. strain red) were very close hits to *T. urticae* strain green and appeared to reflect variation at sites subject to reverse transcription misincorporations (see modification index methods below). This is consistent with *T. urticae* being a very common greenhouse pest in Colorado [71], strongly indicating that *T. urticae* was the primary or sole contributing *Tetranychus* species. Filtered reads were then mapped back to a final reference set of *T. urticae* strain green mitochondrial tRNA genes (reference set determined below) requiring that any mapped read represented ≥60% coverage of a reference tRNA and had an e-value < 1e-3.

### tRNA coordinate determination

The degenerate nature of mitochondrial tRNAs in Acariformes has resulted in historical difficulties in finding and defining the coordinates (i.e. the start and stop position) of tRNA genes. In order to construct a reference set of tRNAs based on tRNA transcription data, the “closest” function in bedtools (ver. 2.27.1, [72]) was used to assign reads to a gene based on any amount of overlap with a previously annotated tRNAs. Reads that had overlap with more than one tRNA gene were assigned to the gene with which the greatest overlap occurred. The most frequent start and stop position for each gene were used as the final reference coordinates for mapping (supp. Table 1). Additional transcribed stem-loops were found with the same protocol by identifying all reads that did not overlap with a previously annotated tRNA gene. Final references for all genes can be found in supp. Table 5.

### Folding predictions

Folding prediction analysis was done with RNAFold (ver. 2.4.11, [73]), using the most frequent YAMAT sequence for each gene with a 100% nucleotide identity to the genomic reference. The majority of models presented in Fig. 2 represent the maximum free energy (MFE) folding prediction. See supp. Table 3 for alternative folding predictions (centroid model). Because YAMAT-seq adapters ligate to the four unpaired bases on the 3′of RNAs, folding predictions were constrained to have an unpaired CCA and discriminator base and pairing involving the first base (position 1 and *n* - 4). For antisense tRNA-Arg, antisense tRNA-Cys, tRNA-Ile, antisense tRNA-Phe, tRNA-Pro, tRNA-Ser1, tRNA-Thr, antisense tRNA-Val, and the D-loop sequence, the MFE model had an internal budge with a weakly base-paired structure. In these cases, the centroid or the MFE model is presented after excluding these weakly supported internal base pairs (Fig. 2). The folding models for tRNA-Asp, tRNA-His, and tRNA-Trp were based on Chen et al., 2014 except that weak base pairing in putative T-arms was excluded.

## Acknowledgements

We would like to thank Laurence Drouard and Thalia Salinas-Giegé for assistance with developing plant (mite) mitochondrial isolation techniques. We also thank the Colorado State University greenhouse staff for plant care and apologize for any past complaints about pest outbreaks. This work was supported by graduate fellowships from the National Science Foundation (DGE-1450032) and the Embassy of France in the United States (Chateaubriand Fellowship), as well as start-up funding from Colorado State University and a grant from the National Institutes of Health (R01 GM118046).

