## Supplemental Figure 1 for "Hopeful monsters: Unintended sequencing of famously malformed mite mitochondrial tRNAs reveals widespread expression and processing of sense-antisense pairs"

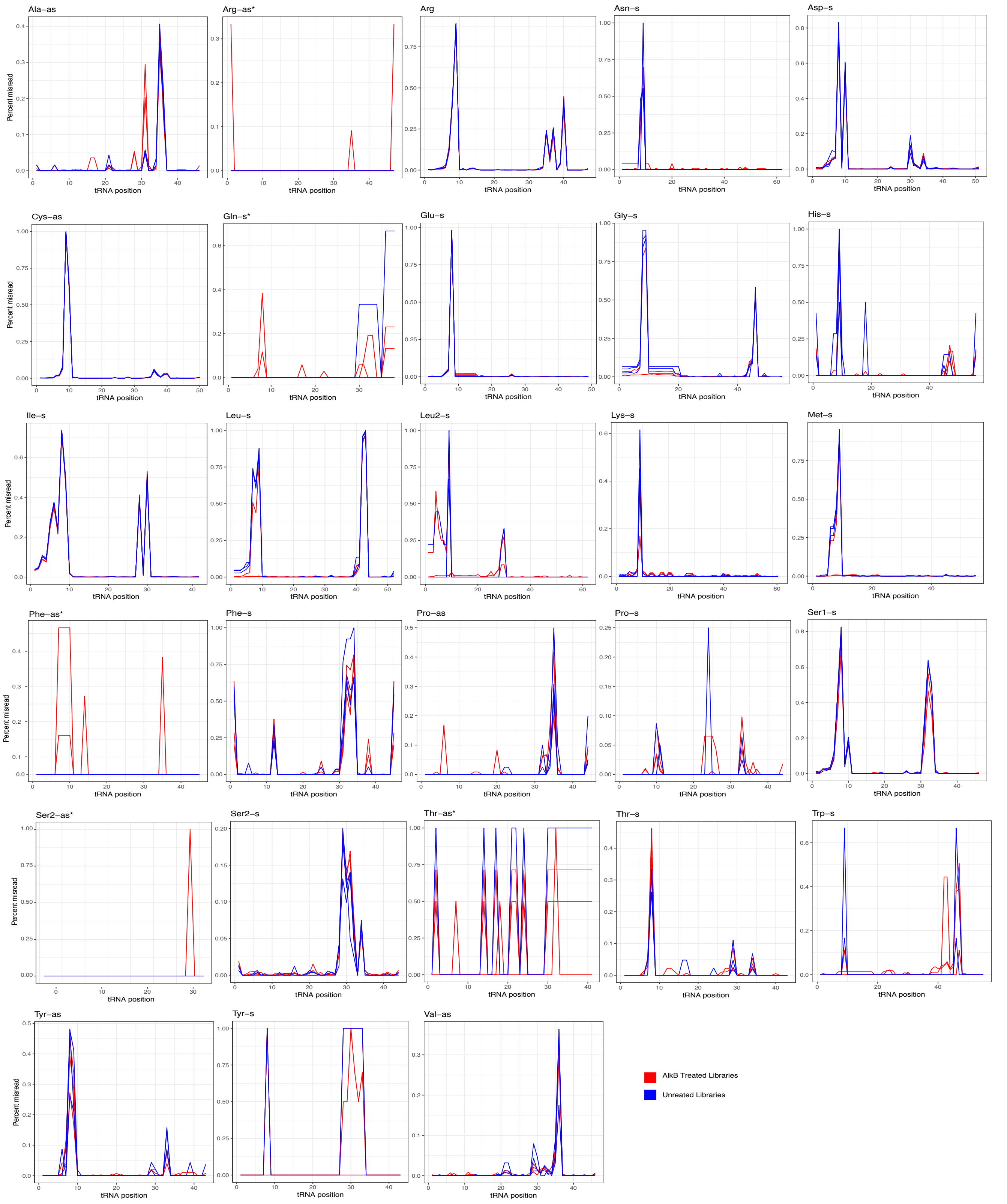


S1 Fig.| Modification indexes of *T. urticae* mitochondrial tRNAs. Percentage of reads that differed from genomic reference for each position of each mt-tRNA. tRNAs marked with a * represent tRNAs with low coverage, reducing the confidence in the modification index.
